# Subjugating Translational Inhibition in Response to Ribosomal Protein Insufficiency by a Herpesvirus Ribosome-Associated Protein

**DOI:** 10.1101/2020.10.11.334607

**Authors:** Elizabeth I. Vink, John Andrews, Carol Duffy, Ian Mohr

## Abstract

In addition to being required for protein synthesis, ribosomes and ribosomal proteins (RPs) also regulate mRNA translation in uninfected and virus-infected cells. By individually depleting 85 RPs using RNAi, we found overall protein synthesis in uninfected primary fibroblasts was more sensitive to RP-depletion than those infected with herpes simplex virus-1 (HSV-1). Although representative RP-depletion (uL3, uS4, uL5) inhibited protein synthesis in cells infected with other DNA viruses, HSV-1-infected cell protein synthesis unexpectedly endured and required a single virus-encoded gene product, VP22. During individual RP-insufficiency, VP22-expressing HSV-1 replicated better than a VP22-deficient variant. Furthermore, VP22 cosedimented with ribosomes and polyribosomes in infected cells. This identifies VP22 as a virus-encoded, polyribosome-associated protein that compensates for RP-insufficiency to support viral protein synthesis and replication. Moreover, it reveals an unanticipated class of virus-encoded, ribosome-associated effectors that reduce the dependence of protein synthesis upon RPs and broadly support translation during physiological stress such as infection.

## INTRODUCTION

Comprised of 40S and 60S subunits assembled from four rRNAs and 79 ribosome proteins (RPs) conserved among eukaryotes, mammalian 80S ribosomes are immensely complex ribonucleoprotein machines that shape the proteome vital for all cells and their intracellular parasites. Originally regarded as invariant, homogenous components universally present in all cells charged exclusively with decoding mRNA, accumulating evidence suggests that the protein composition of ribosomes is actually heterogeneous and this can serve to regulate gene expression. Several *S. cerevisiae* RPs are encoded by duplicated genes where some paralogs are functionally distinct and differentially responsive to stress [1, 2]. RP abundance in multicellular organisms varies between cell and tissue type with up to 25% of all RPs exhibiting tissue-specific expression, and many RPs accumulate in sub-stoichiometric amounts in mammalian cells [3, 4]. Furthermore, translation of a discrete subset of mRNAs encoded by the murine HOX genes is reliant upon a specific RP, eL38 (RPL38) to support mouse skeletal development [5]. Thus, together with changes in ribosome abundance, heterogeneous ribosome pools may influence gene expression by engendering differential selectivity of translated transcripts. Besides influencing differentiation and development, RP haploinsufficiency causes ribosomopathies, a diverse group of human clinical disorders resulting in tissue specific developmental pathologies where ribosome dysfunction is believed to play an underlying role [6]. Finally, RP levels impact cellular homeostasis and the response to physiological stress, including virus infection [2, 7, 8].

To subvert ongoing host cell mRNA translation and enforce production of the polypeptides required for their own reproduction, viruses deploy numerous strategies to access cellular ribosomes [8, 9]. One tactic exposes differential requirements for RPs in translating viral versus cellular mRNAs. Cis-acting internal ribosome entry sites (IRES) in RNA virus genomes (Cricket Paralysis virus, encephalomyelocarditis virus, poliovirus, hepatitis C virus) are reliant upon eS25 (RPS25), which is largely dispensable for host cell cap-dependent mRNA translation [10, 11]. Similarly, different RNA virus IRESs (foot and mouth disease virus, classical swine fever virus, Seneca valley virus) rely upon eL13 (RPL13) [12]. In contrast, flaviviruses exploit P1 and P2 (RPLP1 and RPLP2) components of the 60S stalk substructure for translation of mRNA encoding the envelope protein and Vesicular Stomatitis Virus is dependent upon eL40 (RPL40) for translation of its capped viral mRNAs. [13-15] Finally, DNA viruses can also target RPs as Vaccinia virus modifies the RP Receptor for Activated C Kinase 1 (RACK1) to effectively customize the host ribosome and enhance viral protein synthesis. [16] However, precisely how RPs might influence the lifecycle of other DNA viruses, including those in the medically important herpesvirus family remains unknown.

Although virus reproduction is largely suppressed in human peripheral neurons latently-infected with Herpes Simplex Virus-1 (HSV-1), the virus episodically exits latency and accesses epithelial tissue where productive replication and virus shedding ensue. [17] During its productive growth program, HSV-1 impairs ongoing host mRNA translation by accelerating mRNA decay and restricting new host transcription to allow viral m^7^G-capped, polyadenylated mRNAs to better compete for limiting translational components and in part to stifle host anti-viral responses. [18-21] Multiple, independent virus-encoded functions conspire to constitutively activate the cellular cap-dependent translation machinery by inactivating the translational repressor 4E-BP1 and promoting assembly of the eIF4F multi-subunit initiation factor complex that recruits 40S ribosomes to the mRNA capped 5’-terminus. [22-25] Although the capacity of HSV-1 to stimulate the cellular machinery needed for translation initiation is well characterized, how the ribosome itself might control infected-cell protein synthesis and the relative reliance of viral protein synthesis upon specific RPs remains unknown.

To evaluate how changes in RP levels influence translation in HSV-1-infected primary human fibroblasts, an siRNA screen targeting individual RPs was performed and infected cell protein synthesis and virus reproduction were measured. In marked contrast to uninfected cells, overall protein synthesis in HSV1-infected cells was unexpectedly found to be far less sensitive to specific RP-insufficiency induced by individually depleting 85 core ribosome proteins and paralogs. While depletion of uL3 (RPL3), uS4 (RPS9), or uL5 (RPL11) inhibited protein synthesis in both uninfected cells and in cells infected with other DNA viruses, global protein synthesis in HSV-1-infected cells persisted. Continued protein synthesis and enforced productive replication during uL3, uS4, or uL5 insufficiency was dependent upon a single HSV-1 gene product, VP22 in the presence or absence of a functioning virus endoribonuclease *vhs* allele. Together these results identify VP22 as a viral factor that compensates for RP-insufficiency to support viral protein synthesis and replication. Moreover, they define VP22 as an unanticipated virus-encoded effector with potential to broadly support virus protein synthesis and replication during RP-insufficiency. Rather than relying on specific or modified RPs to enforce viral mRNA translation, VP22 may broadly support viral mRNA translation in different cell types and / or physiological conditions despite variations in the host cell RP portfolio.

## RESULTS

### Translation in HSV-1-infected cells resists RP-depletion

To determine how RP insufficiency impacts mRNA translation in HSV1-infected cells, an RNAi screen targeting 85 RPs and RP paralogs was executed (Fig 1A). Following transfection of control, non-silencing or individual RP-specific siRNAs into primary human fibroblasts (NHDFs), cell number was quantified 48h later to first assess viability and proliferation. While some minor variation was observed, the vast majority of siRNAs did not significantly alter cell number (Table S1). Thus, under these conditions, depletion of an individual host RP in primary NHDFs did not detectably interfere with cellular viability. To evaluate how RP-depletion influenced ongoing protein synthesis in mock-infected versus HSV-1-infected NHDFs that were treated with control, non-silencing or individual RP siRNA, nascent polypeptides were metabolically radiolabeled with ^35^S-amino acids at 15 hpi and protein synthesis rates were quantified by measuring acid insoluble radioactivity. Results from depleting individual RPs were averaged and plotted relative to translation rates in non-silencing, control siRNA-treated cells to compare how RP-depletion in general influenced protein synthesis in mock versus HSV-1-infected NHDFs. When visualized as aggregate data sets, RP-depletion reduced translation rates by approximately 42% in mock-infected cells, indicating as expected that protein synthesis in normal primary fibroblasts was strongly dependent upon RP sufficiency (Fig 1B). By contrast, protein synthesis was reduced by only 14% upon RP-depletion in NHDFs infected with HSV-1 (Fig. 1B). Individual RP siRNAs were next grouped according to their impact on protein synthesis in mock- and HSV-1-infected cells. 60 siRNAs reduced translation rates at least 30%, 24 had minimal impact on translation, and one increased translation rates at least 30% in mock-infected cells (Fig 1C, Fig S1, Table S2). However, only 23 RP siRNAs reduced translation rates by at least 30%, while 55 RP siRNAs showed only a minor reduction and 7 siRNAs increased infected cell protein synthesis by at least 30% in HSV1-infected NHDFs. Taken together, these results demonstrate that: i) protein synthesis in HSV-1 infected NHDFs is more resistant to RP-insufficiency in response to individual RP-depletion by RNAi than mock-infected cells; and ii) a greater number of individual RP-siRNAs inhibited protein synthesis in mock-infected cells than HSV1-infected cells. This reveals a surprising, fundamental difference in how normal vs. HSV1-infected primary fibroblasts responds to ribosome protein sufficiency. Moreover, it reveals how specific RP-depletion differentially impacts translation in uninfected-vs HSV-1-infected cells.

**Figure 1.**
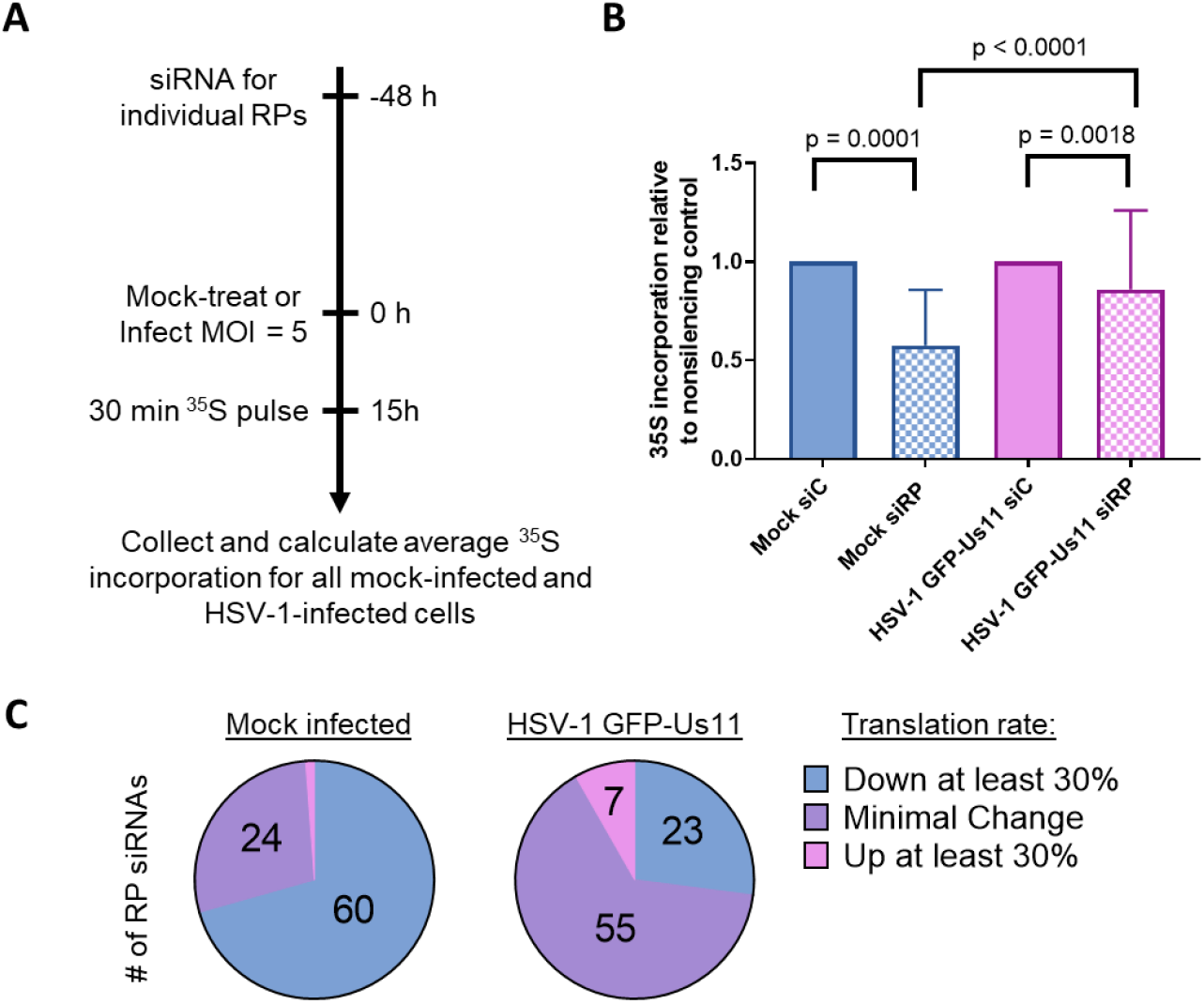
Translation in HSV-1-infected cells resists RP depletion. **A)** Screening strategy to investigate how depleting individual host cell ribosome proteins impacts translation in mock- and HSV-1-infected NHDFs. **B)** NHDFs treated with non-silencing control siRNA or siRNA targeting each individual RP were mock-treated or infected with HSV-1 at an MOI of 5. Approximately 15 hpi cells were metabolically pulse labeled with ^35^S labeled amino acids for 30 min and protein synthesis was quantified by counting acid insoluble radioactivity in liquid scintillant. An aggregate data set was compiled by averaging together^35^S incorporation for all mock-infected cells and compared to an aggregate data set compiled by averaging together ^35^S incorporation in all HSV-1-infected cells Error bars = standard deviation (S.D.) **C)** Number of RP siRNAs that inhibit translation, increase translation, or have only minor impacts on translation in mock- and HSV-1-infected cells.

### HSV-1 enforces protein synthesis when uL3, uS4, or uL5 are limiting

To gain mechanistic insight on how individual RPs might differentially impact translation in mock vs HSV-1-infected NHDFs, three representative RPs (60S proteins uL3, uL5 ; 40S protein uS4) whose depletion resulted in a large decrease in global protein synthesis in uninfected cells, but had minimal impact in HSV-1-infected cells were selected for further study. First, metabolically labeled proteins isolated from NHDFs treated with control, non-silencing or individual RP-targeted siRNA that were mock-infected or infected with HSV-1 (Fig. 2A) were compared. The efficacy with which HSV-1 infection impairs ongoing host cell protein synthesis and directs virus protein production is evident upon comparing mock vs HSV-1-infected cultures treated with non-silencing, control siRNA (*Fig. 2A, compare lane 1 to 3, 5 to 7, 9 to 11*). Compared to control, non-silencing siRNA-treated cultures, uL3, uS4, or uL5 -depletion did not detectably alter the qualitative spectrum of protein synthesis in either infected or uninfected NHDFs (Fig 2A). However, although uL3, uS4, or uL5 -depletion reduced protein synthesis by approximately 45% in mock-infected cells, it did not significantly alter translation rates in HSV-1-infected cells effectively validating the initial screen results (Fig. 2A, B). To evaluate how uL3, uS4, or uL5 -depletion influenced virus reproduction and spread, infectious virus production in a multi cycle growth experiment was quantified by plaque assay. Depletion of uS4 resulted in a small 4-fold reduction in yield relative to non-silencing control treated cells, while depletion of uL3 or uL5 did not significantly impact replication (Fig. 2C). These results reinforce that RP-depleted NHDFs are not only viable, but further demonstrate that primary fibroblasts depleted for uL3, uS4, or uL5 still support robust HSV-1 productive replication with at best only extremely limited interference detected. Moreover, it indicates that, depletion of an individual host RP in primary NHDFs isn’t simply interfering with cellular viability or capacity to support a complex developmental gene expression program represented here by HSV-1 reproduction.

**Figure 2.**
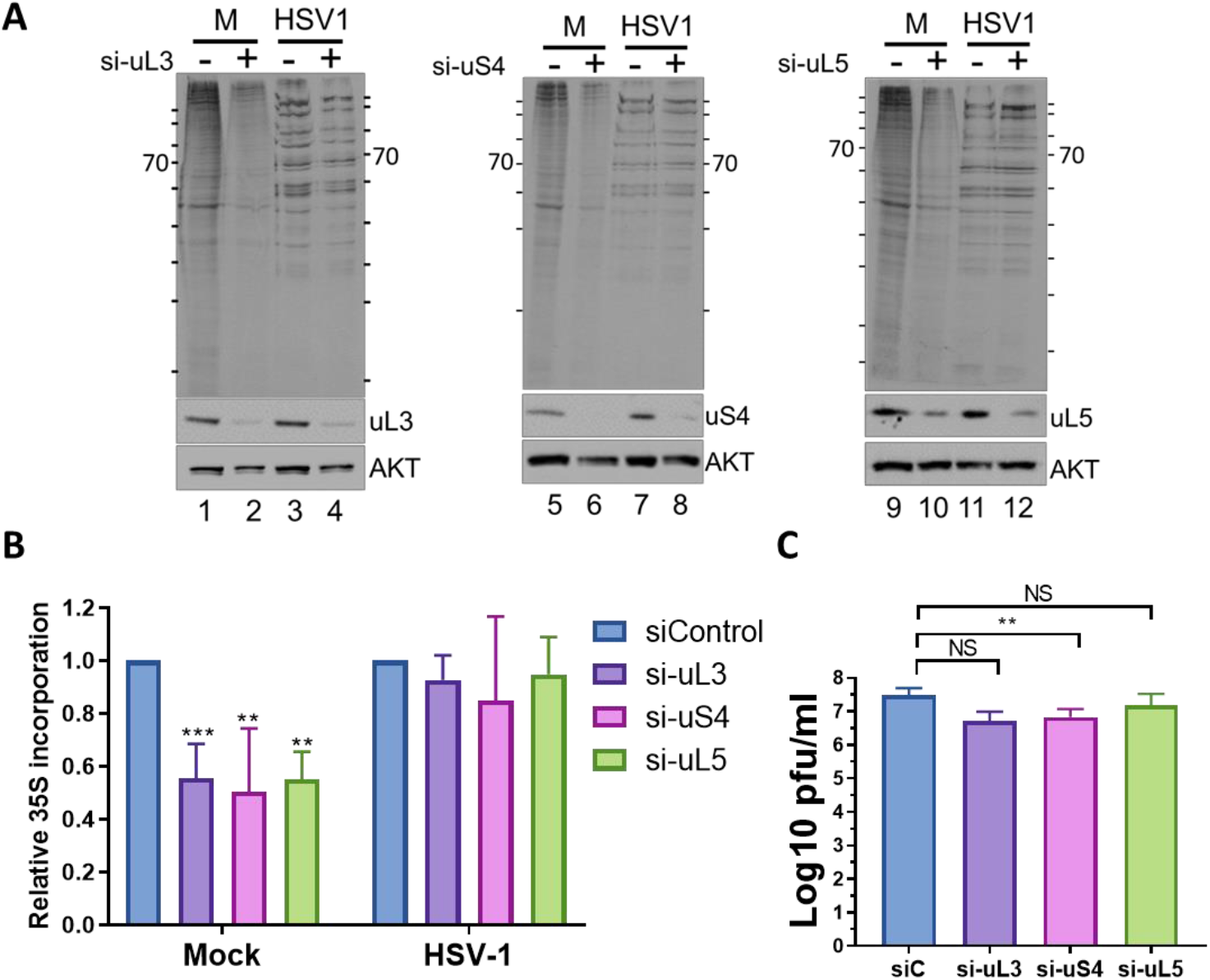
HSV-1 enforces protein synthesis when uL3, uS4, and uL5 are limiting. **A)** NHDFs treated with non-silencing control (siC) or siRNA specific for uL3, uS4, or uL5 were mock-treated or infected with HSV-1 (MOI = 5). Approximately 15 hpi, cells were metabolically pulse labeled with ^35^S AA for 30 min. Total protein was fractionated by SDS-PAGE and the dried gel exposed to X-ray film. **B)** As in (A) except that protein synthesis was quantified by counting acid insoluble radioactivity in liquid scintillant. Results are normalized to nonsilencing control siRNA. N = 5. **C)** NHDFs treated with the indicated siRNA were infected with HSV-1 GFP-Us11 (MOI= 2 x10^−4)^. At 48 hpi, cell-free lysates were prepared by freeze-thawing and infectious virus produced was quantified by plaque assay. N = 3. Error bars = S.D. * = P < 0.05, ** = P < 0.01, *** = P < 0.001.

### Translation in cells infected with either of two different DNA viruses remains sensitive to uL3, uS4, or uL5 - depletion

To determine if the relative resistance of protein synthesis to RP-depletion is a generalized property of primary cells to DNA virus infection or is unique to HSV-1, how RP-depletion influenced ongoing translation in cells infected with a related human herpesvirus that replicates in the nucleus (human cytomegalovirus, HCMV) was examined. Unlike HSV-1, which impairs ongoing host protein synthesis and reprograms the host cell to almost exclusively translate viral mRNA, the betaherpesvirus HCMV sculpts the host translatome to stimulate production of both viral and host proteins which enhances infectious particle production [26]. These two disparate strategies of viral translation control, where one virus (HSV-1) impairs host protein synthesis while the other (HCMV) does not could have differential requirements for individual RPs. Alternately, insensitivity of translation to RP-depletion could represent a conserved property of herpesvirus-infected cells. Analysis of metabolically-labeled proteins in NHDFs treated with control, non-silencing or RP-targeted siRNAs revealed that uL3, uS4, or uL5 -depletion did not detectably alter the pattern of ongoing protein synthesis observed following fractionation of total protein by SDS-PAGE (Fig. 3A). However, unlike HSV-1 infected cells (Fig. 2A,B), RP-depletion reduced protein synthesis both in mock-infected and HCMV-infected NHDFs by approximately 50% (Fig 3B). As interfering with ribosome biogenesis triggered by HCMV-infection regulates IFNB1 production [27], the capacity of a small molecule inhibitor of JAK, which blocks IFNB1 production and signaling, to potentiate the effects of RP-depletion on protein synthesis in HCMV-infected cells was evaluated. Protein synthesis in HCMV-infected NHDFs remained sensitive to RP-depletion, and was not detectably ameliorated by JAK inhibitor treatment (Fig S2), indicating that sensitivity of HCMV-infected cell protein synthesis to RP-depletion is independent of JAK signaling and INFB1 induction. Thus, in marked contrast to HSV-1-infected cells, HCMV-infected cell protein synthesis remains similarly sensitive to uL3, uS4, or uL5-depletion as mock-infected primary fibroblasts. While this finding is inconsistent with a shared herpesvirus function conserved between HSV-1 and HCMV that confers resistance to RP-depletion, it remains possible that resistance of protein synthesis to RP-depletion results in part from the strong impairment of host protein synthesis in HSV-1-infected cells.

**Figure 3.**
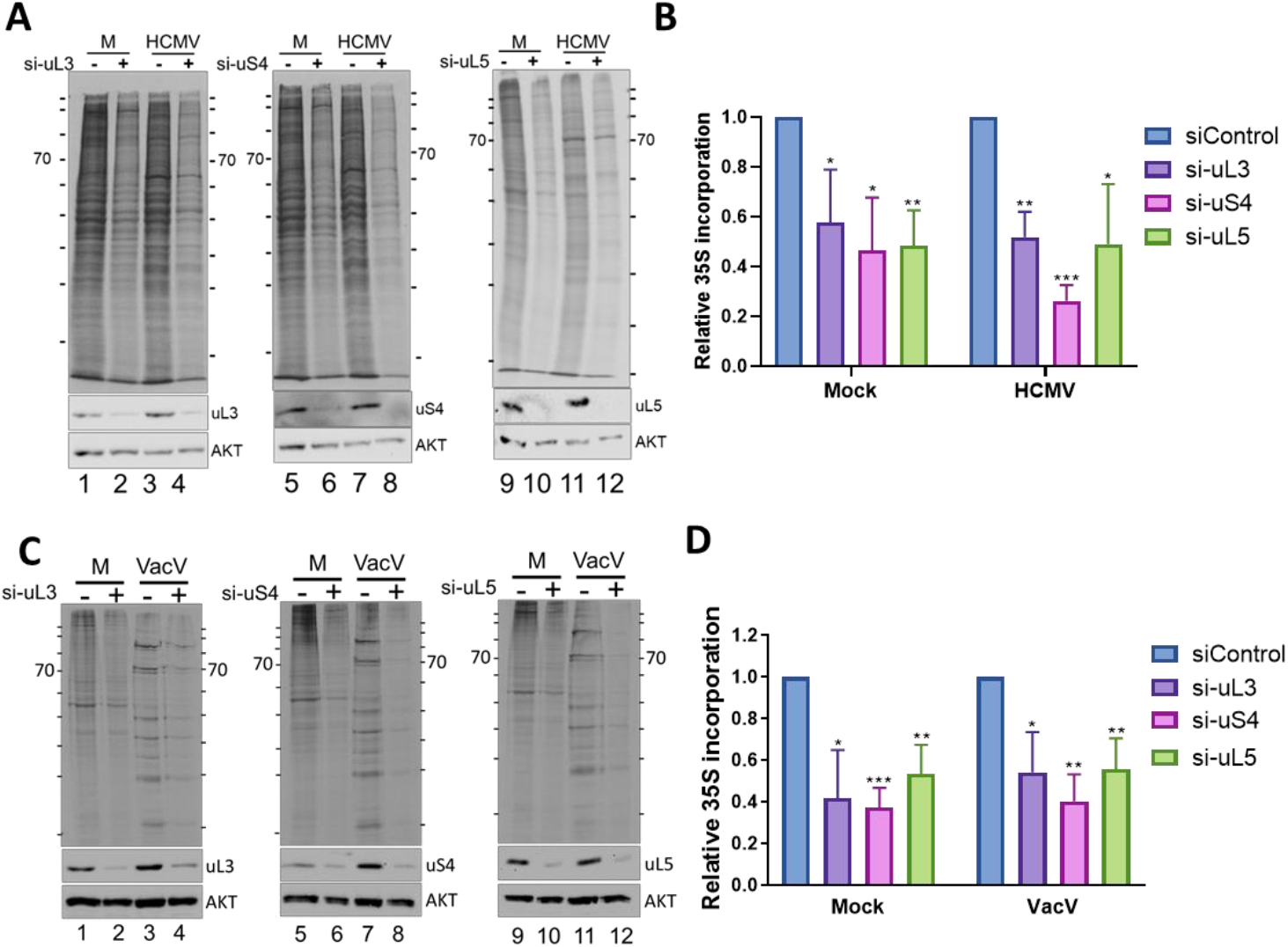
Translation in HCMV- and VACV-infected cells is sensitive to uL3, uS4, and uL5 depletion. **A)** NHDFs treated with non-silencing control (-) or siRNA specific for uL3, uS4, or uL5 were mock-treated or infected with HCMV (MOI = 3). Approximately 72 hpi, cells were metabolically pulse-labeled with ^35^S AA for 30 min. Total protein was harvested, fractionated by SDS-PAGE, and analyzed by autoradiography and immunoblotting with the indicated antibodies. **B)** As in **A**, except protein synthesis was quantified by counting acid insoluble radioactivity in liquid scintillant. Results are normalized to non-silencing control (siC). N = 4. **C)** As in A except NHDFs were mock-treated or infected with vaccinia virus (MOI = 5). At 15 hpi, cells were metabolically pulse-labeled for 30 min and analyzed as in A. **D)** Protein synthesis in samples described in C was quantified as in B. Results are normalized to non-silencing control. N = 4 Error bars = S.D. * = P < 0.05, ** = P < 0.01, *** = P < 0.001.

To investigate whether host shut-off imposed by an unrelated DNA virus might influence responses to RP-depletion, the impact of uL3, uS4, or uL5 - depletion upon protein synthesis in cells infected with a poxvirus that replicates in the cytoplasm (Vaccinia virus, VACV) was considered next. Like HSV1, VACV both produces capped, polyadenylated mRNA and accelerates global mRNA decay to inhibit host cell protein synthesis [28]. Comparing the pattern of metabolically labeled proteins isolated from NHDFs treated with control, non-silencing or individual RP-targeted siRNA that were mock-infected or infected with VACV indicates that VACV infection impairs ongoing host cell protein synthesis and reprograms host ribosomes to produce predominately virus-encoded polypeptides (*Fig. 3C, compare lanes 1 to 3, 5 to 7, 9 to 11*). However, compared to cultures treated with control, non-silencing siRNA, protein synthesis in mock-infected and VACV-infected cells were similarly reduced in a statistically significant manner by uL3, uS4, or uL5 -depletion (Fig 3D). Thus, even though both HSV-1 and VACV potently suppress ongoing host protein synthesis via a related mechanism involving accelerated mRNA decay mediated by virus-encoded factors, only protein synthesis in HSV-1-infected cells is resistant to uL3, uS4, or uL5-depletion. By contrast, protein synthesis in VACV-infected primary fibroblasts is as sensitive to RP-depletion as mock-infected cells. These results establish that the resistance of infected cell protein synthesis to uL3, uS4, or uL5-depletion is not a universal property of DNA virus-infected cells. Instead, the resistance of infected cell protein synthesis to RP-depletion is selectively observed in HSV-1-infected cells and is consistent with the possible involvement of one or more HSV-1 specific gene products.

### Productive replication during uL3, uS4, or uL5 -insufficiency is dependent upon the HSV1 late protein VP22

To better delineate the HSV-1 genetic determinant(s) required to enforce protein synthesis during RP-insufficiency, the penetrance of this phenotype at discrete kinetic phases within the virus productive growth cycle were first established. Quantification of protein synthesis in metabolically-labeled uL3, uS4, or uL5-depleted NHDFs infected with HSV1 or mock-infected revealed both were similarly reduced by approximately 50% after 4h (Fig. 4A). While protein synthesis in mock-infected primary fibroblasts consistently remains sensitive to RP-depletion, ongoing translation in HSV-1-infected NHDFs becomes insensitive to uL3, uS4, or uL5-depletion by 15 hpi (Fig. 2B). This indicated that resistance of HSV-1-infected cell protein synthesis to RP insufficiency developed or was acquired over the course of the reproductive lifecycle. Although both uninfected and infected cell protein synthesis were similarly sensitive to uL3, uS4, or uL5-depletion early in the virus reproductive lifecycle, only infected cell protein synthesis was insensitive to RP-depletion at later times. This suggested that one or more late virus-encoded gene products might render infected cell protein synthesis resistant to RP-depletion. To test this possibility, HSV-1-infected NHDFs were treated with phosphonoaceitic acid (PAA), which inhibits viral DNA synthesis and thereby prevents HSV-1 late gene expression together with subsequent advancement into the late phase of the reproductive cycle. Figure 4B demonstrates that translation in PAA-treated, HSV-1 infected cells was reduced by approximately 60% upon uL3 or uS4-depletion and 35% by uL5-depletion. This indicates that a process dependent on viral DNA replication or late viral gene expression is required to support infected cell protein synthesis during uL3, uS4, or uL5 insufficiency.

**Figure 4.**
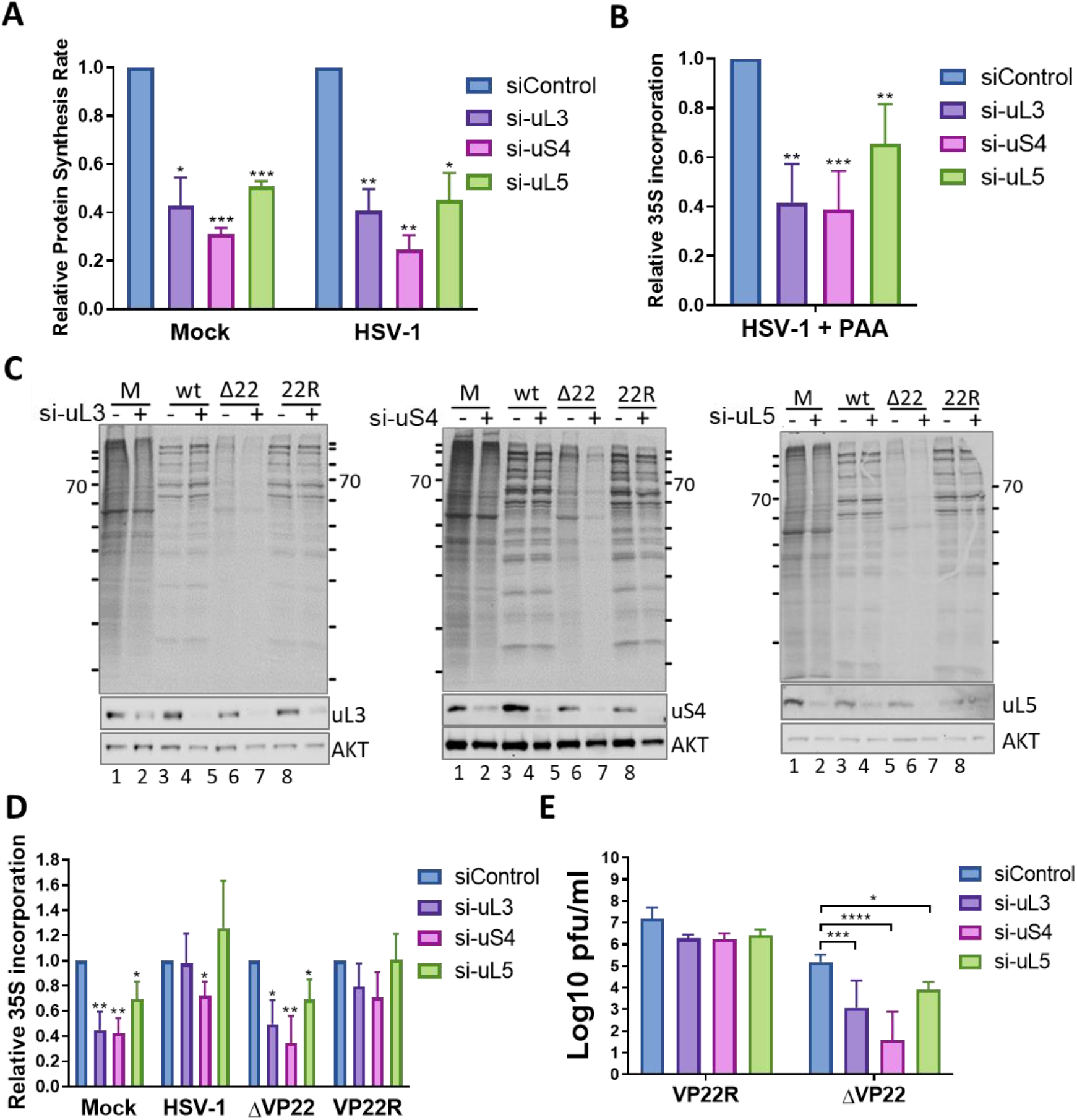
HSV-1 late protein VP22 enforces protein synthesis to support productive replication when uL3, uS4, and uL5 are depleted. **A)** NHDFs treated with non-silencing control (siC) or siRNA specific for uL3, uS4, or uL5 were mock-treated or infected with HSV-1 (MOI = 5). Approximately 4 hpi, cells were metabolically pulse-labeled with ^35^S AA for 30 min. Total protein was isolated and protein synthesis quantified by counting acid insoluble radioactivity in liquid scintillant Results are normalized to non-silencing control siRNA. N = 5, **B)** As in A except cells were treated with PAA to block viral DNA replication and late gene expression and cells were metabolically pulse labeled at 15 hpi. **C)** NHDFs treated with non-silencing control siRNA or siRNA specific for uL3, uS4, or uL5 were mock-treated or infected with wild type (wt), VP22-deficient HSV1 (ΔV22), or VP22-repair HSV1 (22R) (MOI = 5). Approximately 15 hpi, cells were metabolically pulse labeled as in (A). Total protein was isolated, fractionated by SDS-PAGE and the fixed, dried gel exposed to X-ray film. D) Protein synthesis in samples described in (C) was quantified by counting acid insoluble radioactivity in liquid scintillant. N = 4. E) NHDFs were treated with siRNAs as indicated in (C) and infected with HSV-1 ΔVP22 Repair (VP22R) or HSV-1 ΔVP22 (ΔVP22) (MOI = 2 * 10^−4^). At 48 hpi, cell-free lysate was prepared by freeze-thawing and infectious virus produced was quantified by plaque assay. N = 4 Error bars = S.D. * = P < 0.05, ** = P < 0.01 *** = P < 0.001.

Among the approximately 56 late viral gene products encoded by HSV-1 that might render infected cell protein synthesis insensitive to RP-depletion, the RNA binding protein VP22 emerged as an attractive potential candidate. Prior studies of VP22 had established its role in controlling translation of late viral mRNAs. [29-32] To determine if infected cell protein synthesis during RP-insufficiency was dependent upon VP22, protein synthesis in primary cells infected with a VP22-deletion mutant virus (HSV1-ΔVP22) was compared to an otherwise isogenic repaired virus where the wild-type VP22 allele was restored into its native position (HSV-1-VP22R) [33]. Analysis of metabolically labeled polypeptides produced by NHDFs infected with either WT HSV-1 or HSV-1-ΔVP22R by SDS-PAGE showed that both effectively impaired ongoing host protein synthesis and enforced HSV1 protein synthesis (Fig 4C, compare lanes 1 to 3 and 7). Compared to mock-infected NHDFs, protein synthesis in WT HSV-1 or HSV-1-VP22R-infected cells was largely insensitive to uL3, uS4, or uL5 -depletion (Fig 4C, D). While uL3 and uL5 knockdown did not have a statistically significant impact on translation rates, uS4 knockdown reduced translation rates a modest 28% in WT HSV1 infected cells, but did not have a statistically significant impact in HSV-1 VP22R infected cells (Fig. 4D).

In contrast to viruses carrying a functional VP22 allele, radiolabeled polypeptides produced by NHDFs infected with HSV1-ΔVP22 exhibited an alternate protein expression pattern (Fig 4C) and less overall protein synthesis (Fig. 4D) consistent with earlier published reports [29, 31]. Surprisingly, global translation rates in HSV-1 ΔVP22-infected cells remained markedly sensitive to RP-depletion (Fig 4C, lanes 5, 6 in each panel). uL3, uS4, or uL5-depletion reduced protein synthesis in in HSV-1 ΔVP22-infected NHDFs by 50%, 65%, and 31% respectively (Fig. 4D). The extent to which RP-depletion reduced protein synthesis in in HSV-1 ΔVP22-infected cells was similar to sensitivities observed in mock-infected NHDFs where uL3 or uS4-depletion inhibited translation rates by approximately 65%, whereas uL5-depletion reduced translation rates 30% (Fig. 4D).

To determine whether productive growth of the VP22-deletion virus was hypersensitive to RP-depletion, the replication and spread of HSV-1 ΔVP22 was compared to HSV-1 VP22R. NHDFs, treated with siRNAs targeting uL3, uS4, or uL5, or a non-silencing control were infected with HSV-1-ΔVP22 or HSV-1-VP22R and infectious virus yield was quantified by plaque assay. Depletion of uL3, uS4, or uL5 resulted in respective reductions of 25, 779, or 19-fold in HSV-1-ΔVP22 yield relative to control conditions, whereas knockdown of the three RPs individually did not have a statistically significant impact on HSV-1-VP22R productive replication (Fig 4E). Thus, in the absence of VP22, productive replication and infectious virus production are hyper-sensitive to uL3, uS4, or uL5 insufficiency. Taken together, these results establish that i) protein synthesis in HSV1-infected cells; and ii) HSV-1 productive growth and infectious virus production during uL3, uS4, or uL5 insufficiency are all dependent upon the VP22 late gene product.

### Surmounting RP-insufficiency and regulating VHS are genetically separable VP22 functions

VP22 is a multifunctional protein that binds mRNA and promotes translation at late time points during infection [30, 32]. In addition, VP22 regulates gene expression in infected cells through its physical interaction with the viral endoribonuclease *vhs* [32, 34], While recombinant viruses deficient in VP22 exhibit a cell-type specific reduction in global protein synthesis, VP22-deficient viruses accumulate secondary suppressor mutations within *vhs* upon passaging, establishing a genetic interaction between these two HSV-1-encoded functions [31, 35]. Furthermore, these secondary *vhs* mutations, which reportedly either limit mRNA decay or relieve a *vhs*-dependent block to nuclear mRNA export, restore overall protein synthesis to near wild-type (WT) levels. [31, 34] To verify and quantify how *vhs* influenced VP22-enforced mRNA translation, total protein synthesis was compared in NHDFs infected with a recombinant HSV-1 harboring a VHS endoribonuclease-inactivating point mutation in the presence (HSV1 VHS D213N) or absence of VP22 (HSV-1 ΔVP22-/VHS D213N). [31] While protein synthesis in cells infected with HSV-1 VP22R, which is phenotypically indistinguishable from WT with respect to protein synthesis (Fig 4) or *vhs* D213N were similar, protein synthesis in HSV-1 ΔVP22-infected cells was reduced by approximately two-fold (Fig 5A). Introduction of the VHS D213N allele into a ΔVP22 genetic background increased protein synthesis in infected NHDFs between 2-3 fold (Fig 5A) in agreement with published reports. [31]

**Figure 5.**
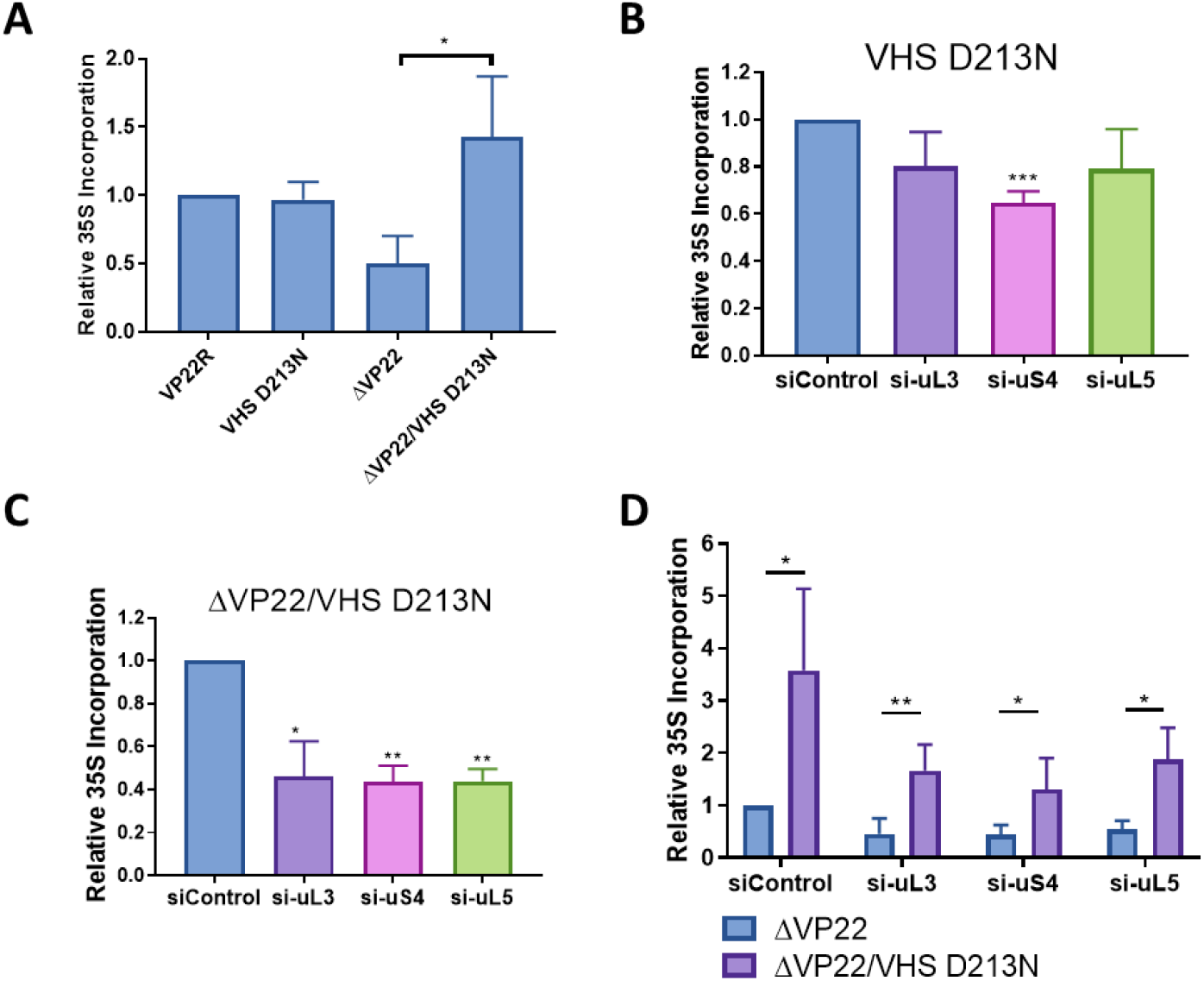
Regulating VHS endoribonuclease activity and subverting RP-insufficiency are independent VP22 functions. **A**) NHDFs were infected with HSV-1 ΔVP22-repair, VHS endoribonuclease mutant HSV-1 VHS D213N, VP22 deletion mutant HSV-1 ΔVP22, or VHS VP22 double mutant HSV-1 ΔVP22/VHS D213N (MOI = 5). At 15 hpi, cells were metabolically pulse-labeled with ^35^S AA for 30 mi. Total protein was isolated and protein synthesis quantified by counting acid insoluble radioactivity in liquid scintillant. N = 4. B) NHDFs treated with non-silencing control or siRNA targeting uL3, uS4, or uL5 were infected with HSV-1 VHS D213N (MOI = 5). At 15 hpi, cells were metabolically radiolabeled with ^35^S AA and protein synthesis quantified as in **A**. N = 4. C) As in **B** except cells were infected with HSV-1 ΔVP22/VHS D213N. N = 3. D) As in **B** except cells were infected with HSV1 ΔVP22 or HSV1 ΔVP22/ VHS D213N N = 4. Error bars = S.D. * = P < 0.05, ** = P < 0.01, *** = P < 0.001.

To evaluate whether *vhs* endoribonuclease activity contributes to protein synthesis during RP insufficiency, NHDFs treated with siRNAs targeting uL3, uS4, or uL5 or non-silencing control, were infected with HSV-1 VHS D213N. Relative to HSV1 VHS D213N-infected, non-silencing control treated NHDFs, depletion of uL3 or uL5 did not result in a statistically significant reduction of protein synthesis, whereas uS4-depletion inhibited translation by approximately 40% (Fig 5B). In contrast, depletion of uL3, uS4, or uL5 all reduced translation rates by greater than two-fold in cells infected with doubly-deficient HSV1 ΔVP22/VHS D213N (Fig 5C). This indicates that a secondary *vhs* mutation introduced into a HSV-1 VP22-deficient background does not detectably enable protein synthesis during RP-insufficiency. Direct comparison of protein synthesis in NHDFs infected with either HSV-1 ΔVP22 or double mutant HSV-1 ΔVP22/VHS D213N revealed that while overall levels of protein synthesis were greater with the inactivating VHS allele D213N, infected cell protein synthesis was similarly reduced by approximately two-fold upon uL3, uS4, or uL5 depletion (Fig 5D). Thus, while the VHS D213N allele increased the magnitude of protein synthesis in cells infected with VP22-deficient HSV-1, it did not detectably ameliorate the sensitivity of infected cell protein synthesis to RP-depletion. Taken together, these results indicate that VHS endoribonuclease activity is dispensable for VP22-dependent protein synthesis during RP-insufficiency. Furthermore, the capacity of VP22 to sustain protein synthesis during RP-insufficiency represents an unanticipated activity that is genetically separable from its reported role in regulating VHS.

### VP22 cosediments with polyribosomes

The newly identified capacity of VP22 to enforce translation during RP-insufficiency raised the possibility that VP22 might execute this function in part by targeting ribosomes directly. To determine whether VP22 could associate with ribosomes, cytoplasmic extracts from HSV-1 infected NHDFs were fractionated over a 10-50% sucrose gradient and migration of RNA evaluated by monitoring A254nm in each fraction (Fig 6). Total protein isolated from each fraction was analyzed by SDS-PAGE followed by immunoblotting using antibodies that specifically recognize individual ribosomal proteins or virus-encoded polypeptides. As expected, uL3, uS4, or uL5 cosedimented with free 40S (uS4) and 60S (uL3, uL5) subunits, were most abundant in fractions corresponding to the 80S monosome peak, and also detected throughout light and heavy polyribosome fractions, the latter indicating the most heavily translated mRNAs (Fig 6). Significantly, VP22 was found to cosediment with RPs both in fractions enriched in ribosome subunits, 80S monosomes and light and heavy polysomes (Fig 6). ICP27, which was previously found to associate with PABPC1 and polyribosomes [36-39] provides a positive control for a known HSV1-encoded, polyribosome ribosome-associated protein sedimented in a similar manner to VP22. By contrast, HSV-1 ICP0 protein migrated more towards the top of the gradient, was not detected in gradient fractions enriched for light or heavy polyribosomes, and served as a negative control. Association of ICP0, with numerous potential protein partners, including translation elongation factor EF-1δ [[40]], could account for migration of a subpopulation with the 80S ribosome peak. These results establish that HSV-1 VP22 cosediments with ribosomes and polyribosomes in infected cells and identify VP22 as a virus-encoded protein that can associate with ribosomes and polyribosomes.

**Figure 6.**
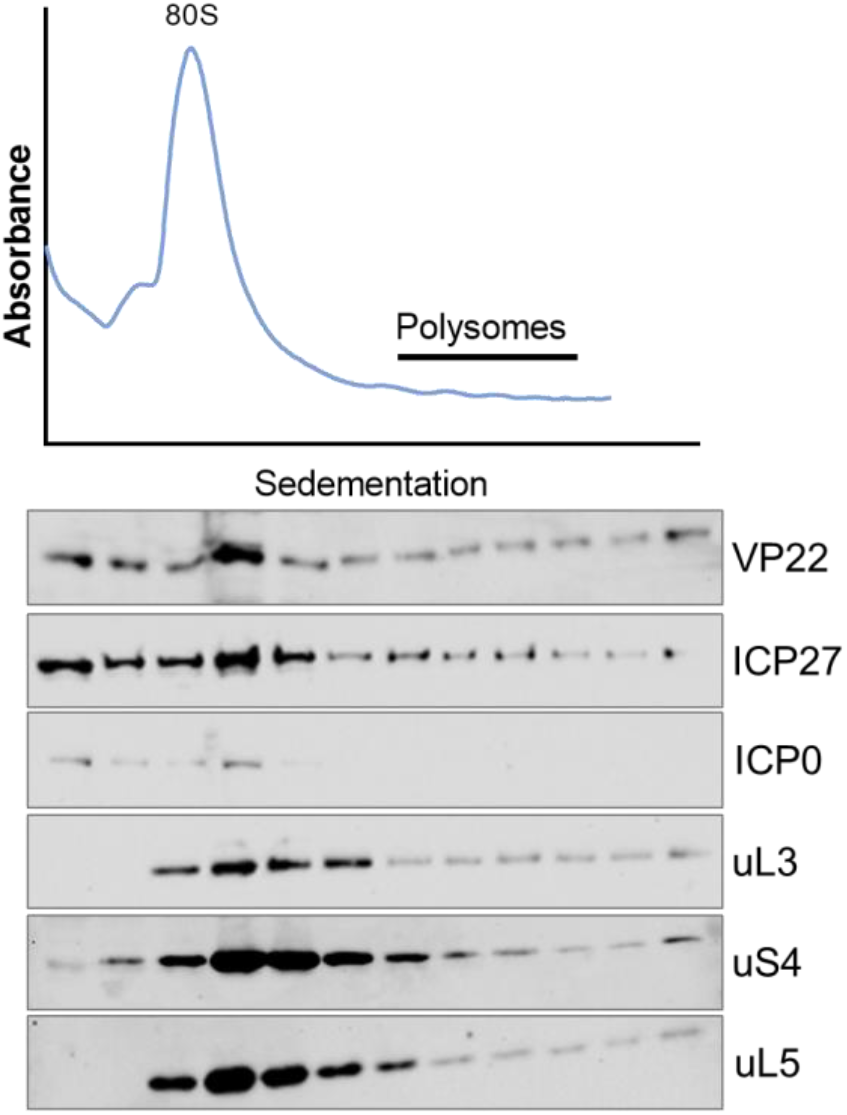
VP22 cosediments with ribosomes and polyribosomes in HSV1-infected cells. NHDFs were infected with phenotypically wild type HSV-1 (MOI = 5). At 15 hpi, cytoplasmic lysate was prepared and fractionated over a 10-50% sucrose gradient. Absorbance at 254 nm was recorded. Total protein in individual fractions was precipitated with TCA, separated by SDS-PAGE, and analyzed by immunoblotting using the indicated antibodies.

## DISCUSSION

Rather than functioning as an invariant ribonucleoprotein complex devoted solely to decoding mRNA, ribosomes are now recognized as dynamic, heterogeneous components that actively regulate mRNA translation. Altering ribosome protein content provides a powerful mechanism for cells to tune gene expression by modifying ribosome selectivity for discrete mRNA populations during normal cell growth and development [1, 4, 41-43] and in response to changing physiological conditions [44], including stress responses like virus infection. Whereas differential RP requirements have been reported to support translation of some virus-encoded mRNAs [11, 14-16, 45-48], how altered ribosome protein content impacts HSV1 infection remains largely unknown. Through an siRNA screen targeting 85 core ribosome proteins and paralogs, we demonstrate here that protein synthesis in normal, uninfected cells is reduced by RP-insufficiency, while protein synthesis in HSV-1-infected cells proceeded despite individual knockdown of 55 RPs. In addition, the majority of 23 RPs required for HSV1-infected cell protein synthesis were required for bulk cellular mRNA translation. In contrast to HSV-1-infected cells, where protein synthesis was enforced despite depletion of individual, representative RPs uL3, uS4, or uL5, protein synthesis in HCMV- and VACV-infected cells was reduced, indicating that translation during RP-insufficiency is a selective property of HSV-1 infected cells. Resistance to RP-insufficiency is acquired late in the virus reproductive cycle, differentiating it from mRNA translation prior to viral DNA replication, and shown to be dependent upon VP22, an HSV1-encoded protein that cosediments with ribosomes and polyribosomes. Besides revealing an unanticipated function for VP22, these results define a new class of virus-encoded effectors that associate with ribosomes and reduce the dependence of infected cell protein synthesis upon host RPs.

While VP22 counteracts the inhibitory effects of RP-insufficiency on translation, the underlying mechanism(s) remains unclear. As a component of the virus particle, VP22 is delivered into the infected cell cytoplasm prior to viral gene expression. It is, however, unable to support protein synthesis during RP-insufficiency until later times, possibly resulting from much greater VP22 accumulation as a late protein or its differential post-translational modification [49, 50]. Alternatively, high-level expression of more abundant virus late mRNAs might impose greater sensitivity to existing, finite RP-levels in acutely-infected cells, rendering their proper translation VP22-dependent. By physically interacting with the viral endoribonuclease VHS, VP22 contributes to the temporal regulation of viral gene expression. However, our genetic analysis demonstrates that VP22 regulation of VHS is distinct and independent from its ability to support protein synthesis during RP-insufficiency, Furthermore, translation in cells infected with vaccinia, an unrelated DNA virus that inhibits host protein synthesis in part by accelerating global mRNA decay, remains sensitive to uL3, uS4, or uL5 -insufficiency. Thus, simply remodeling the cytoplasmic mRNA pool in virus-infected cells is insufficient to enforce protein synthesis during RP-insufficiency. Instead, VP22-dependent protein synthesis during RP insufficiency may require its association with polyribosomes. Similar to ICP27, a viral RNA binding protein that associates with PABC1 [36-38], VP22 is enriched in polysome fractions that contain both large and small ribosomal proteins. The exact determinants recognized by VP22 on ribosomes remain to be defined and could include mRNA, rRNA and/or RPs.

How association of VP22 with ribosomes allows translation during RP-insufficiency, while unknown, could potentially involve altering or remodeling ribosomes by a virus-encoded component. The resulting modified ribosomes could be better suited to support continued translation during physiological stress in response to infection. Conceptually, targeting the ribosome to sustain protein synthesis during physiological stress or unfavorable growth conditions has its foundations in the bacterial stringent response [51-53]. In eukaryotes, ribosomes lacking either specific RPs or the full complement of RPs are present in cells and can impact translation in both the budding yeast S. cerevisiae and mouse embryonic stem cells [4, 54-57]. Altering RP composition by mutations in RP genes results in developmental defects in plants [41] and multicellular animals [58], influences ER stress responses in yeast and mice [59, 60] and RP haploinsufficiency results in diseases termed ribosomopathies in humans that are associated with transcript-specific differences in translational efficiency [43, 61-63]. Conversely, deregulated RPL15 expression and the resulting changes in mRNA translation reportedly promote breast cancer metastasis [64]. RP expression and modification are also differentially regulated by environmental conditions [65-69] and can support mRNA translation and adaptation in response to physiological stress [70-73].

Differential ribosome composition is poised to impact virus infection biology, including balancing host antiviral responses and viral gene expression to influence productive virus replication. As host and viral transcripts can have varying requirements for individual RPs, ribosome composition and heterogeneity can regulate ribosome access and shape the infected cell proteome. Examples of specific RPs or RP modifications largely dispensable for bulk cellular translation but required to support translation of viral transcripts have been identified for both RNA and DNA viruses [10, 15, 16]. While effective at capturing specific ribosomes and possibly assisting viral mRNAs to compete for limiting resources, over reliance upon specific ribosomal proteins for virus mRNA translation, however, may come with an intrinsic vulnerability. Notably, RP levels naturally fluctuate and are responsive to changing physiological conditions [54, 66, 68, 71, 73], and this might adversely impact virus reproduction, Rather than relying upon specific RPs or specialized ribosomes for virus mRNA translation, a contrasting strategy to overcome RP-insufficiency by maximizing plasticity is operative in HSV-1 infected cells. Our results indicate that unlike virus mRNAs whose translation requires one or more specific RPs, HSV1 late mRNA translation is broadly tolerant of individual RP insufficiency. Conceivably, this capacity to overcome RP-insufficiency could bestow an adaptive advantage and better support viral mRNA translation and replication in varied cell types and under different physiological conditions where RP levels might fluctuate.

## METHODS

### Cell Culture, Viruses, and Chemicals

Normal Human Dermal Fibroblasts (NHDFs; purchased from Lonza, Walkersville, MD) were grown in DMEM supplemented with 5% Fetal Bovine Serum. Vero cells (ATCC) were grown in DMEM supplemented with 5% calf serum. HSV-1 GFP-Us11 (Patton strain) was described previously [74]. F-strain viruses HSV1 WT, HSV1 ΔVP22, HSV-1 VP22 repair, HSV1 VHS D213N, and HSV1 49-/VHS D213N are described in [31, 33] and grown in VP22-expressing Vero cells in 15% fetal bovine serum [75]. Vaccinia virus (Western Reserve strain) was handled as described [76] and HCMV AD169 was a gift from Dong Yu and prepared in NHDFs as described in [27]. Pyridone 6/JAK inhibitor I (Millipore 420099-500UG) was dissolved in DMSO and used at a concentration of 10 μM. Phosphonoacetic acid (PAA) was obtained from Sigma (P6909). Cells were pretreated with 300 ug/ml PAA prior to infection, this concentration of PAA was maintained for the remainder of the experiment.

### Antibodies and siRNAs

RPS9 (18215-1-AP) and RPL3 (11005-1-AP) antibodies were purchased from Proteintech. RPL11 (D1P5N), and AKT (9272) were purchased from Cell Signaling Technology. AllStars negative control siRNA was purchased from Qiagen. siRNAs targeting ribosome proteins were purchased as an RNAi Cherry Pick library from Dharmacon. On Target Plus siRNAs (Dharmacon) were reordered for experiments following the screen. To deplete ribosome proteins, NHDFs were transfected with siRNAs using Lipofectamine RNAImax (Invitrogen) according to the manufacturer’s instructions.

uL3 sense 5’ GAUGAAUGCAAGAGGCGUU.

uS4 sense 5’ GAAGCUGAUCGGCGAGUAU.

uL5 sense 5’ UAAAGGUGCGGGAGUAUGA.

### Quantification of Protein synthesis

NHDFs, seeded in a 24-well plate, were treated with siRNAs and infected 48h later. At 15 hpi (HSV1 or VacV) or 72 hpi (HCMV) cells were metabolically pulse labeled with methionine- and cystine-free DMEM supplemented with 70 μCi/ml ^35^S-labelled Met/Cys (EasyTag EXPRESS35S, Perkin Elmer). Total protein was boiled in SDS sample buffer and then either fractionated by SDS PAGE and exposed to film or precipitated in 10% Trichloroacetic acid (TCA) on ice for 30 min as described previous [23]. Acid-insoluble protein was collected onto Whatman GF-C filters which were subsequently washed twice in 10% TCA and twice in 100% ethanol. Radioactivity collected on the filters was quantified by liquid scintillation counting. Each sample was collected in duplicate and technical replicates were averaged. P-values were determined by one sample t-tests.

### Multicycle virus growth assay

NHDFs, seeded into 6-well dishes, were treated with non-silencing control siRNAs or siRNAs targeting uL3, uS4, or uL5. Cells were infected with 100 PFU virus in 0.5 ml of medium per well for 1.5 hours after which the virus inoculum was removed and replaced with fresh medium. Infected cultures were collected 48 hpi, freeze-thawed three times, and infectious virus production was quantified by plaque assay on Vero cells. P-values determined by ANOVA with multiple comparisons.

### Fractionation of RNA and associated proteins by sucrose gradient sedimentation

Polysome analysis was performed as previously described [27]. NHDFs, seeded into a 15 cm dish, were infected with phenotypically wild type VP22 repair HSV-1 (MOI = 5). At 15 hpi, cells were treated with 100 μg/ml cycloheximide (Sigma; C7698) for 10 min and collected in polysome lysis buffer (50 mM KCl, 20 mM Tris HC, pH 7.4, 10 mM MgCl2) containing 1% Triton X-100, 10 U/ml RiboLock RNAse Inhibitor (ThermoFisher Scientific; EO0381) and 100 μg/ml cycloheximide. Lysis was allowed to proceed for 10 min on ice, followed by a 1 min 4° C spin at 14,000 RPM to pellet nuclei and mitochondria. Lysates were then layered onto 10-50% sucrose gradients in thinwall polypropylene ultracentrifuge tubes (Beckman Coulter; 331372). Gradients were then centrifuged at 38,000 RPM for 135 min in a SW41Ti rotor (Beckman Coulter; 331362) at 4°C. A Density Gradient Fractionation System (Brandel; BR-188) was used to generate absorbance profiles by measuring the absorbance of RNA at 254 nm of gradients pumped through a flow cell. Protein from each fraction was precipitated in TCA at -20°C overnight. Precipitate was collected by centrifugation at 14,000 RPM for 30 min at 4°C then washed twice in cold acetone before drying and resuspending in SDS-PAGE sample buffer.

## ACKNOWLEDGEMENTS

We thank members of the Mohr laboratory and Angus Wilson for helpful discussions and Angus Wilson for critically reading the manuscript. This work was supported by National Institutes of Health grants GM056927 and AI073898 to I.M. E.V. was supported in part by an American Cancer Society postdoctoral fellowship (grant PF-16-048-01-MPC).

## Authors contributions

E.V, C.D. and I.M. conceived experiments and interpreted data. E.V. and J.A. performed the experiments. E.V., and I.M. wrote the manuscript and E.V., C.D. and I.M. edited the manuscript.

---

**Supplemental figure 1.**
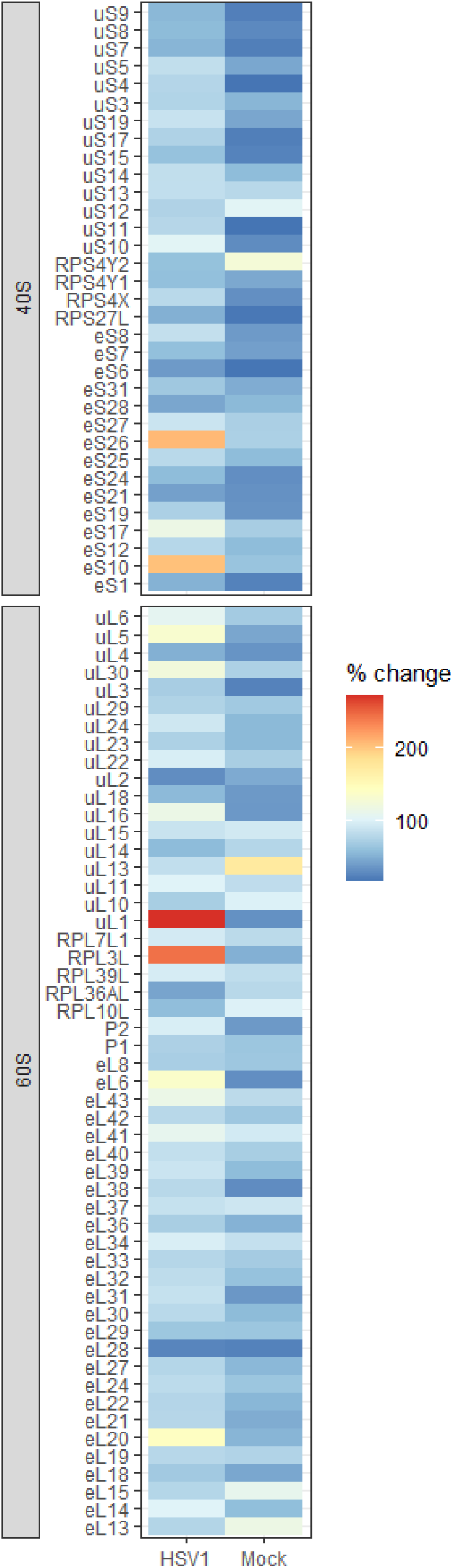
Heatmap summary of RP siRNA-induced changes to global translation rates. Heatmap summary of RNAi screen targeting 85 RPs and RP paralogs as described in Fig 1 and Sup Table 1. Relative protein synthesis rates are presented as the percent change in ^35^S incorporation counts in siRNA-knockdown cells relative to ^35^S incorporation counts in non-silencing, control siRNA-treated cells for both mock-treated, and HSV-1-infected cells.

**Supplemental figure 2.**
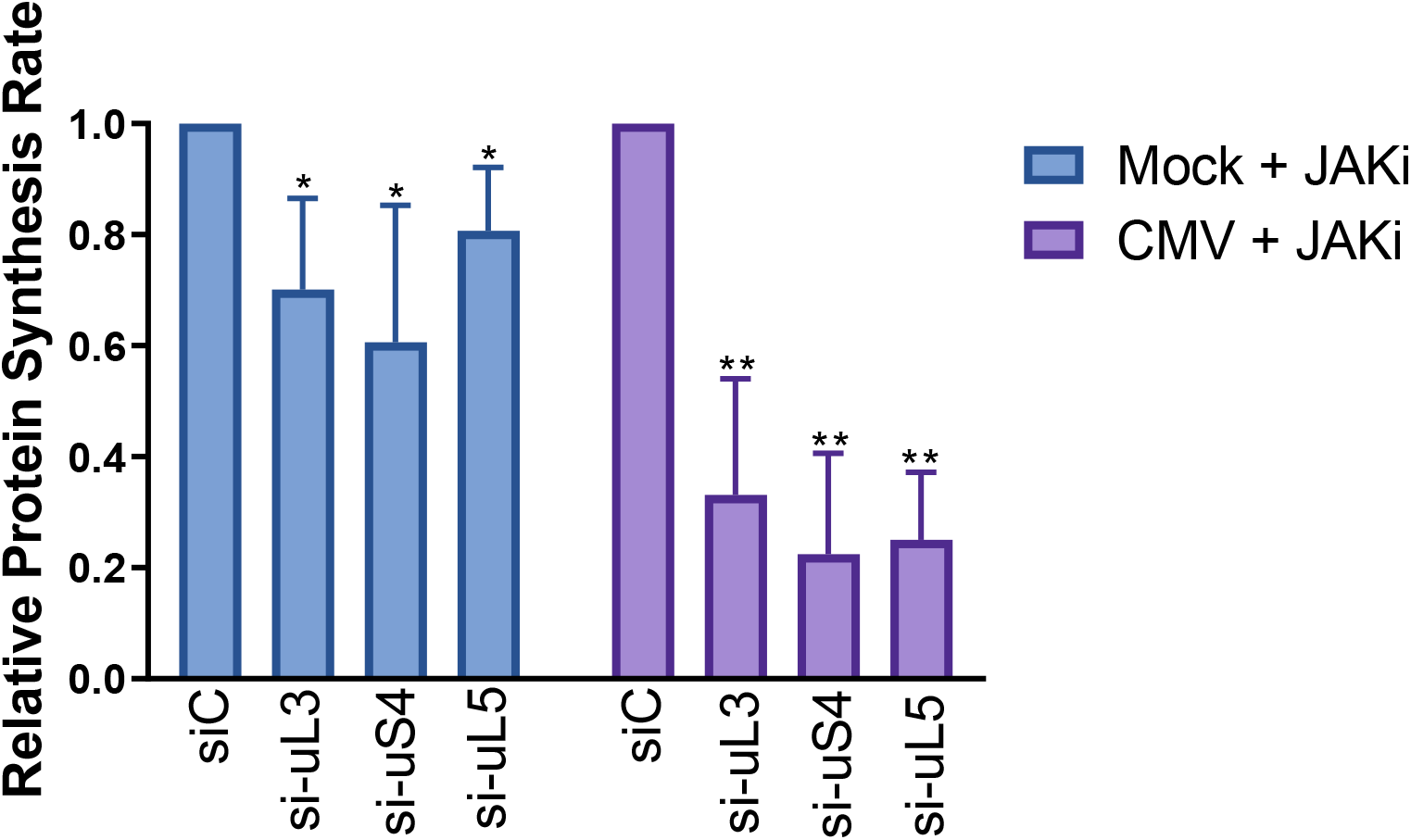
Inhibiting IFN production does not rescue translation in HCMV-infected cells. After treatment with the indicated siRNAs, NHDFs were treated with pyridine 6, a small-molecule JAKi at 10 μM concentration and infected with HCMV (MOI = 3). At 72 hpi, cells were metabolically pulse-labeled for 30 min and protein synthesis was quantified by TCA precipitation followed by counting acid insoluble radioactivity in liquid scintillant. N = probably 4. Error bars = S.D. * = P < 0.05, ** = P < 0.01

**Supplemental Table 1. Impact of RP-depletion on cell number**. Change in cell number due to RP-depletion was determined as follows: approximately 5*10^4^ NHDFs were added to wells in a 24 well plate. The next day, cells were transfected with non-silencing, control siRNA or siRNAs targeting each ribosome protein. Two days later, cell number was counted manually using a hemacytometer and reported as proportion of cell # following RP-knockdown / cells # following non-silencing, control siRNA treatment). N = 3, NS = no statistically significant change in cell number.

**Supplemental Table 2. Impact of RP-depletion on protein synthesis rates**. Results of RNAi screen targeting 85 RPs and RP paralogs as described in Fig 1. Relative protein synthesis rates are presented as ^35^S incorporation counts in siRNA-knockdown cells divided by ^35^S incorporation counts in non-silencing, control siRNA-treated cells for both mock-treated, and HSV-1-infected cells.

